# Belt and braces: two escape ways to maintain the cassette reservoir of large chromosomal integrons

**DOI:** 10.1101/2023.08.31.555669

**Authors:** Egill Richard, Baptiste Darracq, Eloi Littner, Gael Millot, Valentin Conte, Thomas Cokelaer, Jan Engelstädter, Eduardo P.C. Rocha, Didier Mazel, Céline Loot

## Abstract

Integrons are adaptive devices that capture, stockpile, shuffle and express gene cassettes thereby sampling combinatorial phenotypic diversity. Some integrons called sedentary chromosomal integrons (SCIs) can be massive structures containing hundreds of cassettes. Since most of these cassettes are non-expressed, it is not clear how they remain stable over long evolutionary timescales. Recently, it was found that the experimental inversion of the SCI of *Vibrio cholerae* led to a dramatic increase of the cassette excision rate associated to a fitness defect. Here, we question the evolutionary sustainability of this apparently counter selected genetic context through experimental evolution. We find that the integrase is rapidly inactivated and that the inverted SCI can recover its original orientation by homologous recombination between two insertion sequences (ISs) present in the array. These two outcomes of SCI inversion restore the normal growth and prevent the loss of cassettes, enabling SCIs to retain their roles as reservoirs of functions. These results illustrate an interesting interplay between gene orientation, genome rearrangement, bacterial fitness and demonstrate how integrons can benefit from their embedded ISs.

## Introduction

Integrons are ancient genetic structures that play a crucial role in the spread of multidrug resistance in gram-negative bacteria and more generally in bacterial evolution (Cambray et al., 2010; Cury et al., 2016; Mazel, 2006; Neron et al., 2022). These bacterial recombination systems can capture, store and shuffle small mobile elements – cassettes – encoding adaptive functions. They are organized in two parts: a stable platform and a variable array of cassettes (Fig 1a). The stable platform of the integron contains i) the integrase gene (*intI*) under the control of its promoter P_int_, ii) the *attI* integration site and iii) the P_C_ promoter of the cassette that drives the expression of genes encoded in the downstream cassette array. The variable part consists of an array of cassettes, each of which is usually composed of a promoterless gene associated with a recombination site called *attC*. Only the first cassettes (closest to the P_C_ promoter) can be expressed, while the rest represent a low-cost reservoir of valuable functions for the cell (Escudero et al., 2015; Richard et al., 2022a). Many stresses, including the action of certain antibiotics, can trigger the integrase expression (Baharoglu and Mazel, 2014; Guerin et al., 2009). The integrase then catalyzes excision and integration of cassettes in the first position in the array, i.e. at the *attI* site, where they become expressed. The configuration of cassettes that enables the cell to escape stress will be selected. In this way, cassette shuffling enables the bacterium to rapidly screen for all the functions that optimize its survival in a given environment (Escudero et al., 2015).

**FIGURE 1:**
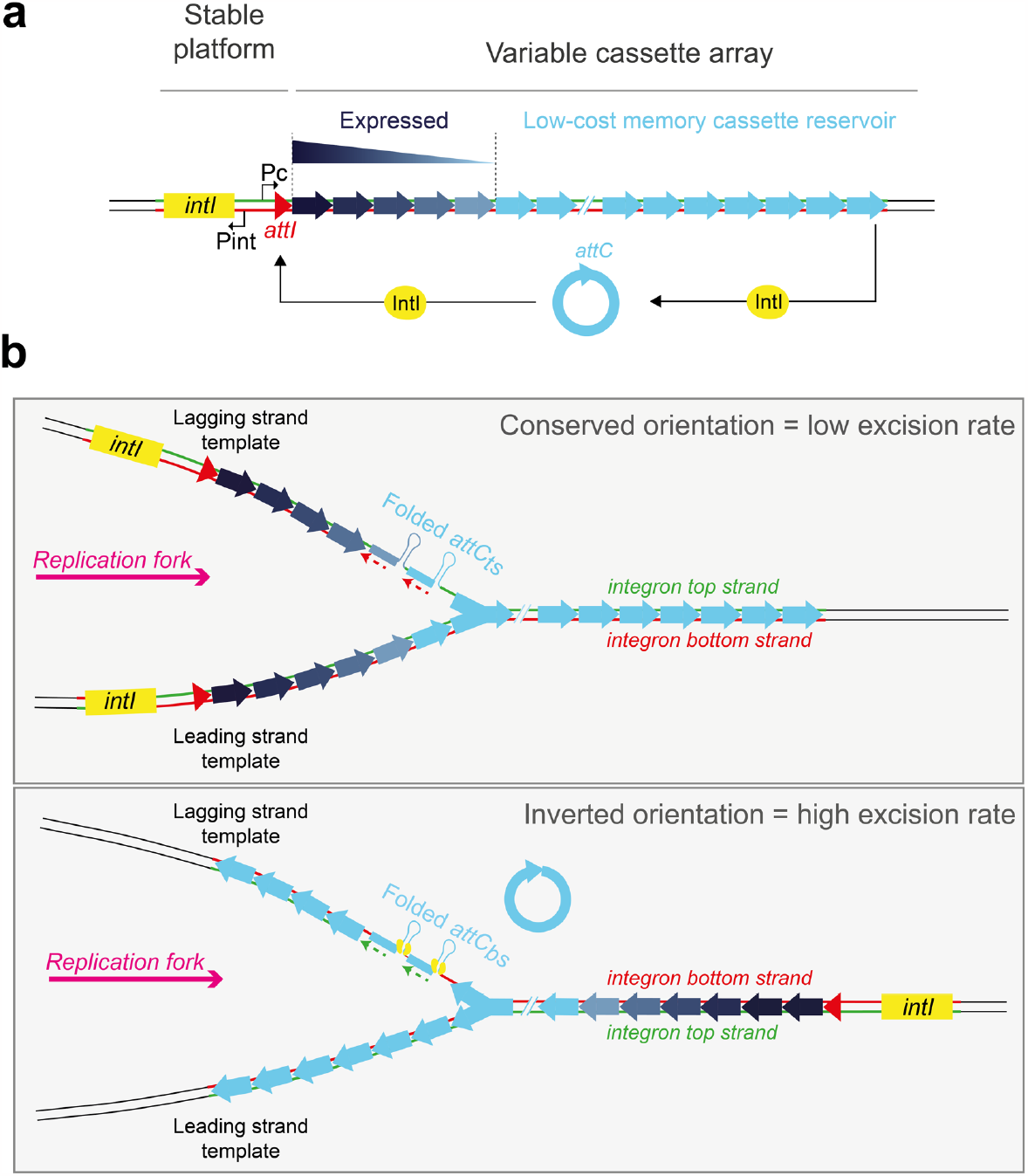
The integron system and the importance of its orientation towards replication. **a**. Schematic representation of the integron. The stable platform consists of a gene coding for the integrase (*intI*, in yellow) and its promoter P_int_, the cassette insertion site *attI* (in red) and the cassette promoter P_C_ driving the expression of the downstream cassettes along an expression gradient. Cassettes are displayed as arrows where the head represent the *attC* site. Their color intensity represents their level of expression from dark blue (highly expressed) to light blue (not expressed). Cassettes can be excised through an intramolecular *attC* × *attC* reaction and be reintegrated in 1^st^ position near the P_C_ promoter through an intermolecular *attC* × *attI* reaction and become expressed. **b**. Mechanistic insight on the issue of integron orientation. Array of cassettes are represented while they are replicated. Their orientation towards the replication fork is indicated by the direction of the arrowhead representing the cassette. In their conserved orientation, SCI recombinogenic bottom strands of *attC* sites (*attC*_*bs*_) are carried by the continuously replicated leading strand template which supposedly limits their structuration. The non-recombinogenic top strands of *attC* sites (*attC*_*ts*_) are carried by the lagging strand template containing stretches of ssDNA (between the Okazaki fragments, dotted lines) which supposedly favors their structuration. In the inverted orientation, recombinogenic *attC*_*bs*_ are carried by the lagging strand template. A more frequent structuration of these *attC* strands is expected to lead to increased binding of the integrase and higher cassette excision rate.

In the context of the ever-increasing pressure imposed by the overuse of antibiotics, integrons are regularly found to encode resistance genes to multiple antibiotics and associated with mobile genetics elements (conjugative plasmids and/or transposons) (Liebert et al., 1999; Stokes and Hall, 1989). These integrons are called Mobile Integrons (MIs) and typically carry small arrays of cassettes probably facilitating their horizontal transfer. However, other integrons are sedentary and located on bacterial chromosomes (sedentary chromosomal integrons, SCIs) (Mazel et al., 1998). SCIs might be the ancestor from which the MIs evolved (Rowe-Magnus et al., 2001). They can easily store dozens of cassettes (up to 301 in *Vibrio vulnificus*), encompassing a substantial fraction of their host genomes (up to 3%) (Cambray et al., 2010; Neron et al., 2022). And yet only the first few cassettes of SCIs are expressed, leaving more than 90% of the cassette array silent. This leads to question: how can such a massive structure that is plastic and mostly not expressed be stable within bacterial genomes over long evolutionary timescales?

In a recent study, we found that the orientation of SCIs towards replication was a crucial factor of their stability (Richard et al., 2022b). The reason for this lies in the form of the recombination substrates of the integron integrase. While this recombinase recognizes the *attI* site in its double-stranded (ds) form through its primary sequence, the *attC* sites are recombined in a single-stranded form (Bouvier et al., 2005; Johansson et al., 2004). Although both the bottom and top strands of an *attC* site can form a secondary structure, only the bottom strand of the *attC* site (*attC*_*bs*_), in its folded form, was found to be recombined by the integrase. This selectivity of the integrase for the bottom strand is essential for the correct orientation of the cassette upon integration at the *attI* site, allowing its expression by the P_C_ promoter (Bouvier et al., 2009; Nivina et al., 2016). It was shown that SCIs are preferentially oriented so that recombinogenic *attC*_*bs*_ are located on the leading strand template where replication is continuous, further decreasing the chance of these *attC*_*bs*_ being structured and recombined (Loot et al., 2017) (Fig1b, top panel). By inverting the SCI of *Vibrio cholerae*, and thus locating the *attC*_*bs*_ on the lagging strand template, we showed that the rate of cassette excision increased dramatically due to a better structuration of the recombinogenic *attC*_*bs*_. This increased cassette excision activity compromises the integrity of the overall structure and also affects the growth of *V. cholerae* in a negative way, mainly due to the increased excision of cassettes encoding functional toxin–antitoxin (TA) systems (Richard et al., 2022b). Here, we assess how this increased integron plasticity due to high cassette excision rate can shape the evolution of SCIs. For that, we use the exceptional plasticity of the inverted SCI of *V. cholerae* as a model to examine the trade-off between evolvability and stability of SCIs through an evolution experiment. We demonstrate that the inverted SCI carrying strain quickly restores its fitness defect and loses its high plasticity in two distinct ways. First, we find that most of the time inactivation of the integrase is rapidly selected, notably through a frameshift induced by an indel occurring in a DNA polymerase slippage hotspot within its coding sequence (CDS). Second, and more strikingly, we find that the array of the inverted SCI can in part recover its original more stable orientation, by homologous recombination between two insertion sequences (ISs) present in the array. In the first way, since integrase functionality is rapidly lost the integrity of the cassette array would be barely affected. Nevertheless, it may be that these cassettes can still be excised and recruited by other MI integrase platforms carried, for example, by incoming plasmids (Rowe-Magnus et al., 2002). In this case, in the long-term, the SCI cassette content could be irretrievably lost. In the second way, the SCI recovers its stability with no impact on its functionality, uncovering an elegant link between large genome rearrangements and evolvability of *V. cholerae*.

## Results

### Experimental evolution reveals rapid loss of the burden of a highly recombinogenic chromosomal integron

In our previous study, we showed that inversion of the SCI of *V. cholerae* led to a dramatic increase of its plasticity associated to a growth defect when the integrase is expressed (Richard et al., 2022b). We reasoned that the inverted SCI could be an ideal model to study the impact of increased cassette dynamics on its evolution. Hence, we set out an evolution experiment using this SCI Inv strain. As described in the Material and Methods section, three biological replicates of the SCI Inv strain containing a low-copy number plasmid expressing the integrase (pSC101::*intIA*) were used to initiate as many separated cultures in a medium that enabled the expression of the integrase (Fig 2a). Each population was propagated daily for 7 days (Cooper, 2018). At the end of each day, D0, D1, D2, D3, D5 and D7 (respectively 0, 10, 20, 30, 50 and 70 generations), cultures were plated. 24 clones were collected, and the growth of each collected clone was measured (Fig 2b). As control, we also performed the same experiment using three biological replicates of strains in which the SCI were re-inverted on site (SCI Reinv) and containing a low-copy number plasmid expressing the integrase (Richard et al., 2022b). Note that for these control strains, we only collected 8 clones for each day of evolution. At D0, we observed a significant growth defect in the SCI Inv strain when the integrase is expressed compared to the corresponding SI Reinv strain (Fig 2b). This severe growth defect observed at D0 in the SCI Inv strain is in line with what we observed in our previous study (Richard et al., 2022b). At each additional day of evolution, we observed a progressive attenuation of the SCI inv growth defect. At D7, this defect disappeared completely, and the growth of both SCI Inv and Reinv strains no longer showed any significant difference. We determined the growth rate of each 24 collected clones for the SCI inv strain (Fig 2c). At D0, all tested clones show a strong growth defect when expressing the integrase. The magnitude of the growth defect, as measured on the impact on the growth rate, is rather severe, with a decrease of the growth rate of about 35%. After 7 days of experimental evolution, this growth defect disappears completely in every clone of each independent population (Fig 2c). We observe that some clones very rapidly lose the growth defect associated with the inversion of the SCI in presence of IntIA, recovering normal growth after only 10 generations (D1). After 30 generations (D3), only 3 of the 24 tested clones retained their growth defect, and 100% of the clones had lost the growth defect after 50 generations (D5). This evolution experiment shows that the cost associated with the inversion of the SCI in presence of the integrase is so high that it is strongly counter-selected, leading to an amelioration after only a few dozen of generations.

**FIGURE 2:**
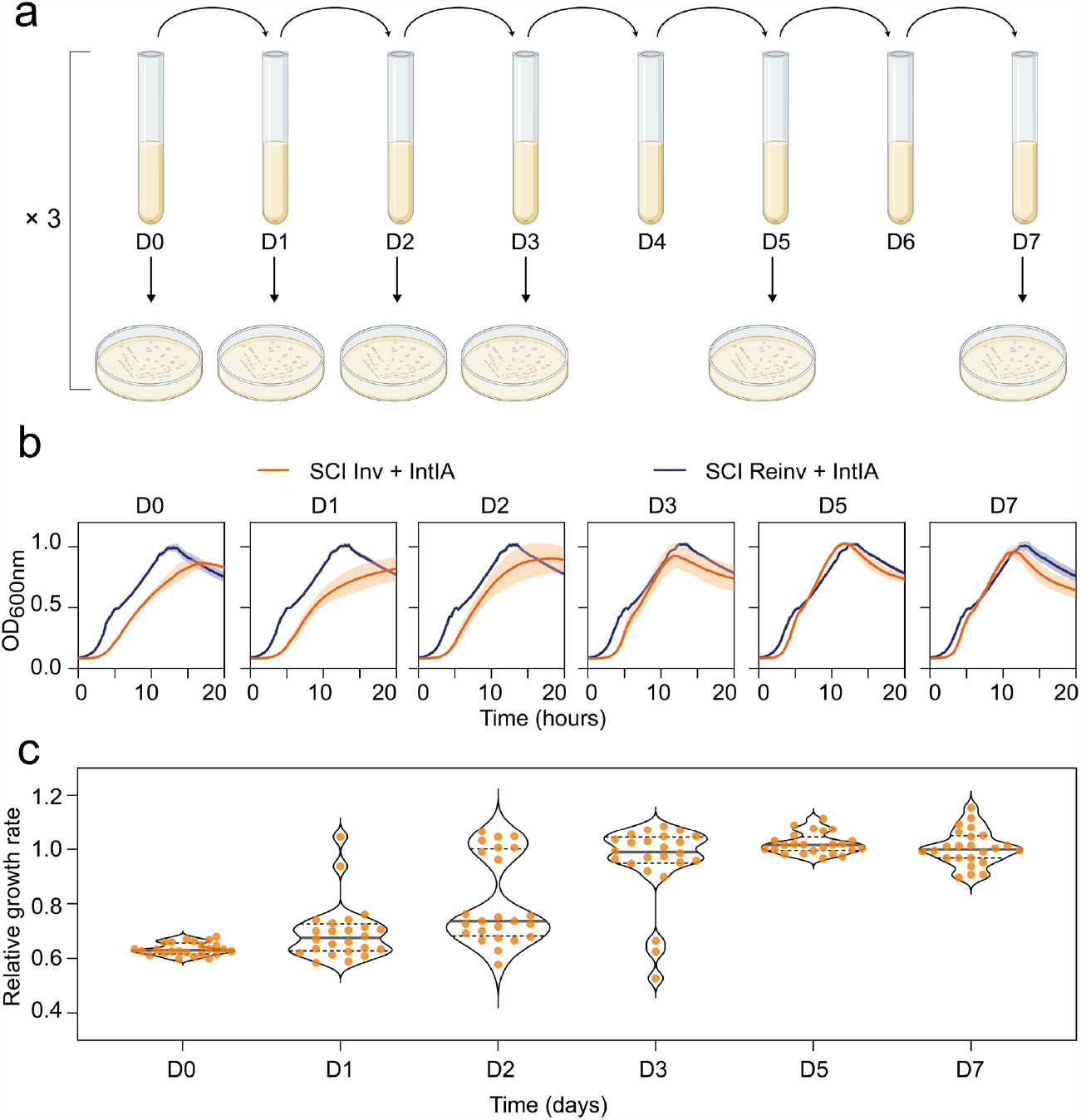
Experimental evolution of the SCI Inv and Reinv strains expressing integrase. **a**. Schematic overview of the evolution experiment. D0, D1, … stands for Day 0, Day 1, and so forth. After each day of growth, cultures were plated. 24 clones and 8 clones were collected for SCI Inv and Reinv strains respectively, both expressing integrase. **b**. Growth curves of the SCI inv (orange lines) and SCI reinv (blue lines) collected clones expressing integrase during the evolution experiment. For each curve, the line corresponds to the mean of the growth of the collected clones, and the shade corresponds to the standard errors at each timepoint. **c**. Distribution of the relative growth rates of the 24 collected clones at each time point of the evolution experiment. The growth rates of each SCI Inv clone for each day are represented as relative values compared to the mean growth rate of the SCI Reinv on the corresponding days. A violin plot is also represented to help visualize the bimodal distributions of the intermediate time points, as well as the median (full line) and quartiles (dotted lines) of those distributions.

### The integron integrase is rapidly inactivated in a highly recombinogenic chromosomal integron

The growth defect of the SCI Inv strain was shown to be associated with high excision activity especially of TA cassettes, so that the quick loss of growth defect was suggestive of a loss of the shuffling activity. To understand the suppression of the growth defect, we first sought to discriminate if the suppressor mutation occurred in the plasmid containing the integrase or if they were of another nature. To achieve this, we cured the plasmid of the 24 clones that were evolved for 70 generations (D7) in the 3 separate populations (as described in the Material and Methods section, and in Fig 3a, top panel). The growth of the 24 SCI Inv clones, before and after the curing of the plasmid containing the integrase, was measured in a medium that enabled the expression of the integrase. The growth of those clones was not found to be different (Fig 3a, bottom panel). Upon retransformation with the plasmid carrying the *intIA* integrase gene, the growth defect was recovered in 21 clones out of 24 (Fig 3a, bottom panel). This showed that the suppressors of the growth defect arose mostly in the plasmid (21/24) but could also be associated to a modification of the chromosome (the remaining 3 clones). We first focused on the more common suppressors, i.e., located on the plasmid. To do that, we extracted the plasmids from the 3 bulk populations evolved for 70 generations, pooled them in equal proportions and sequenced this heterogenous mix of plasmids to a high depth using an Illumina platform. The sequencing of the mix of 6922 bp long plasmid in paired-end 150pb reads resulted in a median coverage of 10,600X. This very high depth of sequencing allowed the calling of variants present at a very low frequency of 0.1%. We found that half of the mutations in the plasmid affected the regulation of the expression of the integrase, mostly through the inactivation of the *araC* gene that is essential to ensure gene expression through the pBAD promoter (see the plasmid scheme Fig 3a, top panel). The rest was found to be in the *intIA* gene (Fig 3b, top panel). Many mutations were present at very low frequencies all along the sequence of the integrase, mostly indels or nonsense mutations. No substitution was found particularly enriched as it could have been expected in the catalytic domains of the integrase like modifications of active-site residues or of the catalytic tyrosine (MacDonald et al., 2006). Instead, indels occurring in DNA homopolymers were found predominant. These indels produce a frameshift thus demolishing the entire integrase protein. One, a stretch of eight “A” starting 57bp after the ATG, was found to gather nearly 40% of all the mutations occurring in the integrase (Fig 3b, top panel). To confirm that this pattern of mutations found in the integrase did not come from a bias in the Illumina sequencing run, we also sequenced the integrase from 43 clones coming from the same 3 populations using classical Sanger sequencing (Fig 3b, middle panel). Even though it offers much less coverage, this approach led to the same general pattern of mutation with an over-representation of indels in the homopolymer of “A” at the beginning of *intIA*. As control, we also extracted the plasmids from the 3 bulk populations evolved for 70 generations from the SCI Reinv strain. As expected, we observed very few mutations in the integrase gene (bottom panel). Thus, we show that integrase mutation is the easiest way to recover from a growth defect associated with overactive recombination, and that this inactivation occurs primarily in a homopolymer motif.

**FIGURE 3:**
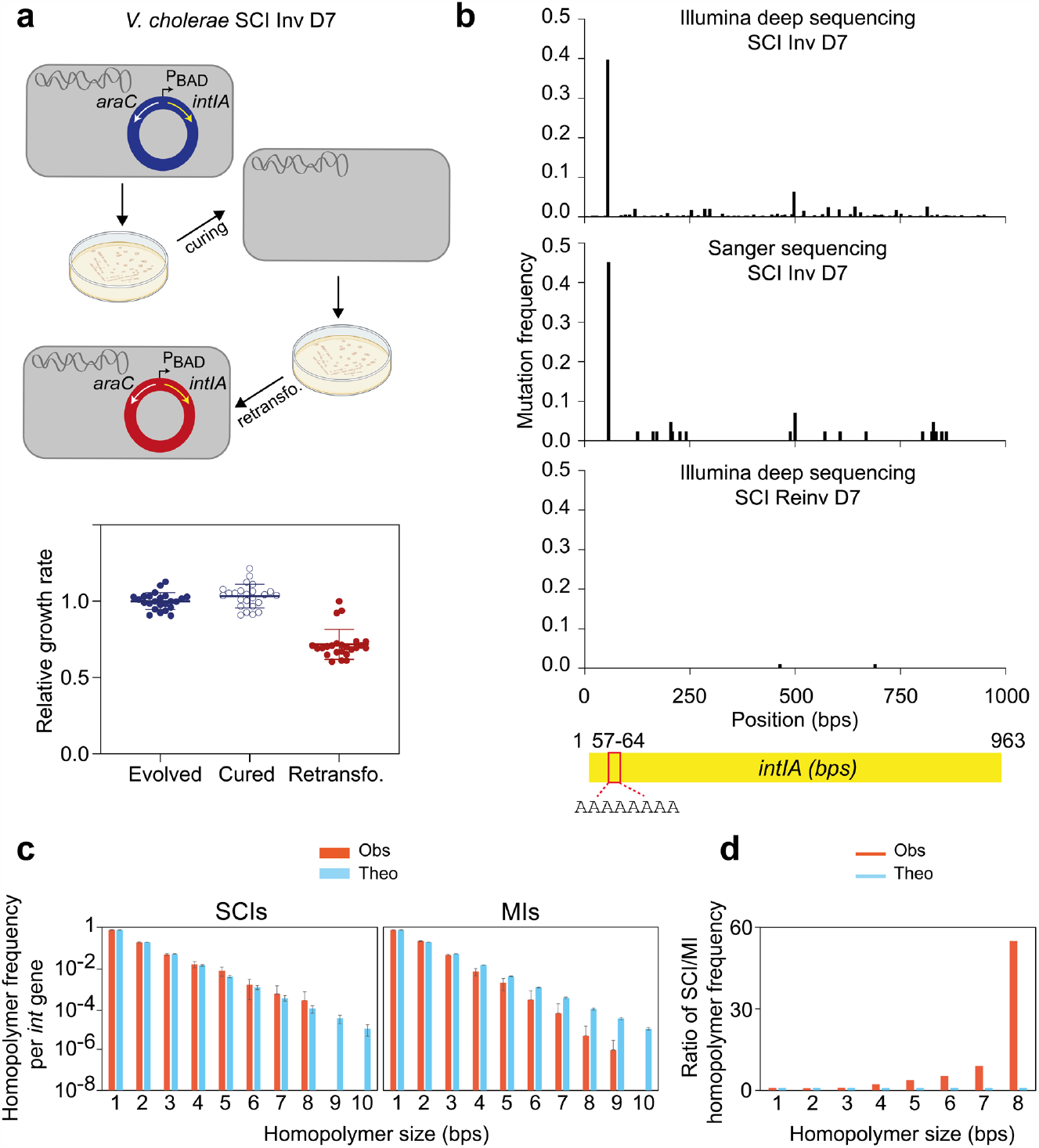
Analysis of the mutation pattern in the *Vibrio cholerae* integron integrase gene. **a**. Schematic overview of the plasmid curing and retransformation procedure and distribution of the growth rates of the 24 clones (before plasmid curing, after plasmid curing and after retransformation of the pSC101::*intIA*). The inducible P_BAD_ promoter, the *araC* repressor gene and the *intIA* integrase genes are respectively represented by black, white and yellow arrows. The growth rates are represented as relative values compared to the mean growth rate of the 24 SCI Inv clones evolved from D7. **b**. Pattern of mutation of the *intIA* gene carried by the pSC101 vector. The x-axis is the nucleotidic position relative to the ATG and the y-axis represents the mutation frequency. Each bar represents a bin of 10 bps that aggregates every type of variants (SNP and indel). The top panel shows results obtained with Illumina deep sequencing approach and the SCI Inv evolved D7 strain (with integrase). The middle panel shows results obtained with Sanger sequencing approach using the SCI Inv evolved D7 strain (with integrase). The bottom panel shows results obtained with Illumina deep sequencing approach using the SCI Reinv evolved D7 strain (with integrase). A schematic view of the integrase gene and the position of the longest homopolymer of the sequence is shown just below the three graphs to help to locate the mutation pattern. **c**. Frequency of the DNA homopolymers for sizes ranging from 1 to 10 within the integrase genes of SCIs (left panel) and MIs (right panel). For each panel, the frequency of the homopolymer sizes observed (obs) in SCI and MI integrase genes is represented in pink and the frequency of the homopolymer sizes theoretically (theo) expected from randomized sequences is represented in blue. The randomized sequences were generated by randomly shuffling the nucleotides of each SCI and MI integrase sequences. **d**. Ratio of the DNA homopolymer frequency for sizes ranging from 1 to 10 within the integrase genes of SCIs compared to the MI ones. The ratio observed (obs) between SCI and MI integrase genes is represented in pink and the ratio theoretically (theo) expected from the randomized sequences is represented in blue.

We then wondered whether the presence of long homopolymers could be a characteristic of integron integrases. For that, we extracted the integrase sequences from the complete integrons identified with IntegronFinder 2.0 in a previous study (Néron et al. 2022). We searched for all the k-mer of nucleotides with k ranging from 1 to 10, termed homopolymers in the following for the sake of simplicity. We then plotted the frequency of the homopolymers in function of their size for SCI (Fig 3c, left panel) and MI integrase sequences (Fig 3c, right panel) and compared each with the expected frequencies obtained from randomized sequences of the same SCI and MI integrase sequences. The distribution of the homopolymer sizes in SCI integrases is almost identical to the randomly simulated distribution. This means that long homopolymers have not been selected to exist in SCI integrases, but still their existence is important because they allow some slippages after SCI inversion. In contrast, the homopolymers in MI integrases are much shorter. A comparison of the homopolymer sizes of SCI versus MI integrases reveals that the 8-nucleotide homopolymers are 47 times less represented in MI integrases than SCI ones, whereas this differential is not observed in the corresponding randomized sequences (Fig 3d). This could reduce the probability of the MI integrases being inactivated by this way.

### Homologous recombination between Insertion Sequences helps maintaining chromosomal integron cassette arrays

Upon retransformation of the evolved clones with the *intIA* plasmid, we identified three SCI Inv clones issued from two different populations, that retained an unaffected growth rate while the others recovered the characteristic growth defect of the SCI Inv strain (Fig 3a). This indicated that the suppressive mutations occurred in the chromosome of these three clones, so we extracted their genomic DNA and queried the presence of these mutations by whole genome sequencing. 100X Illumina sequencing and subsequent variant calling using the sequana pipeline (Cokelaer et al., 2017) showed the absence of any SNPs or indels in these clones, with the exception of SCI in which cassette deletions were found, as we would expect in SCI Inv strains where IntIA was expressed. We therefore asked whether the lack of recovery of growth defects in the three clones could be explained by their cassette content. The use of short reads does not allow an effective assembly of the SCI due to the numerous repeated sequences in that region, so we relied on the coverage of the different cassettes to assess their presence or absence (Fig. 4a). The extensive deletion events that occurred in all three clones did not allow the identification of a specific cassette that was consistently excised in these clones and that could easily explain their phenotype. However, the cassette deletion events observed in all three clones shared the same pattern: a high number of excisions in the first part of the array and almost no excisions in the second part of the array (Fig 4a). Interestingly, the boundary between these high and non-excision parts of the array was identical in all three clones. This boundary involved an insertion sequence (IS) from the IS*3* family that had previously transposed in the CDS of a cassette in the array (Fig 4a). This IS*3* contains two partly overlapping ORFs, as it is characteristic for the IS*3* family (Siguier et al., 2006a; Siguier et al., 2006b), with a total length of 1261 bps. Interestingly, a second copy of this IS*3* is also present in the SCI, inserted in the very last cassette of the array (Fig 4a). The two ISs are inserted in opposite divergent directions. In this configuration, recombination between the two copies of IS*3* would lead to the inversion of the fraction of the cassette array that separates them. We therefore reasoned that in the three clones in which no growth defect could be observed in the presence of the functional IntIA integrase, this event occurred so that most of the array of the SCI was now re-inverted, which could explain the phenotype. We searched for this event performing PCR using a set of four primer couples surrounding the two recombination points: one set for each recombination point and for each orientation of the cassette subset between the two copies of IS*3* (Fig 4b). We observed that in our three clones of interest, the PCR which should be positive in the case of recombination between the two IS*3* copies (r1 and r2) were indeed positive. This is not observed in the 3 clones that we used as controls, which are non-evolved clones (Materials and methods, Fig 4c). Indeed, in these clones, PCR that should be positive in the native inverted configuration (i1 and i2) are indeed positive. This confirms that a recombination event between the two IS*3* did occur in our three clones of interest and led to the inversion of a 79,449 bp segment of the SCI array. Since the recombination involves two IS*3*, we legitimately asked whether the recombination involved the seemingly functional transposases encoded by each copy. First, no target site duplication could be found on each side of the IRL and IRR of both IS*3* copy annotated as such by IS finder (Siguier et al., 2006b), meaning that no recent transposition event involving either copy of the IS*3* occurred. Second, we sequenced each of the previous PCR products and again found no target site duplication in the clones in which the inversion just occurred, indicating that the inversion involved homologous recombination and not transposition events. Note that we found a very weak amplification signal in the PCR r1 and r2 in the non-evolved clones, while no amplification can be observed in the negative controls without DNA template. This means that this recombination event probably occurs at a very low frequency whenever *V. cholerae* is cultivated, even in the absence of integrase, and is selected in condition where it provides an advantage.

**FIGURE 4:**
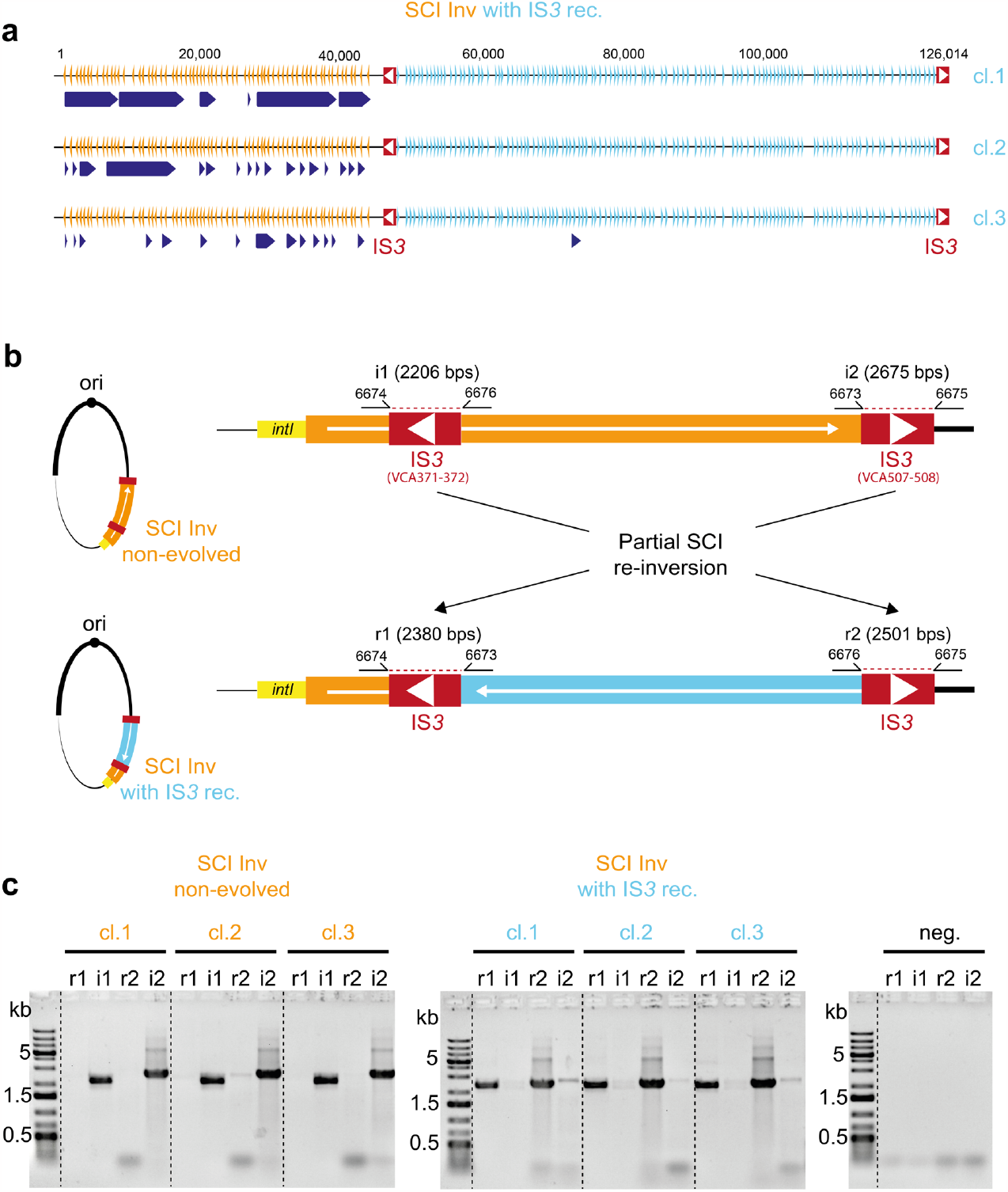
Analysis of the re-inversion events in the array of the *Vibrio cholerae* chromosomal sedentary integron. **a**. Representation of the SCI array of the three clones of interest. The orange triangles show the inverted *attC* sites of each cassette of the array, and the blue ones, the re-inverted *attC* sites. The dark blue arrows represent the deletion events. The red rectangles each with a white arrow show the IS*3* copies. **b**. Schematic representation of the partial SCI re-inversion event at the scale of the chromosome 2 of *V. cholerae* and focus on the two recombination points involved in this re-inversion. The PCR primer couples used to identify the recombination event are represented: i1 and i2 are the couples that should be positive in a non-evolved inverted SCI. r1 and r2 are the couples that should be positive upon recombination between the two IS*3* copies within the inverted SCI (SCI inv with IS*3* rec). **c**. PCR profile of the 6 clones tested for the SCI array configuration. The gels show the PCR results obtained using r1, i1, r2, i2 couple of primers for three non-evolved SCI Inv clones (left panel) as well as for each of the three evolved clones with IS*3* recombination (right panel).

## Discussion

Mathematical models indicate that functional integrase can be stably maintained in population despite substantial fitness costs. This selective advantage arises because cassette shuffling generates a high degree of combinatorial phenotypic diversity, thus enabling the population to respond rapidly to changing selective pressures (Engelstadter et al., 2016). This evolutionary set up would be one of the keys to the success of MIs. Here, concerning large SCIs, we revealed another and more complex layer of regulation which, on the contrary, permit to limit high level of cassette excision thus ensuring the maintenance of large reservoir of cassettes. This regulation occurs thanks to the presence of a pool of special cassettes – the TA cassettes – which become toxic when excised (Jurenas et al., 2022; Richard et al., 2022b). Therefore, all strains with highly recombinogenic SCIs are expected to exhibit a growth defect. This is the case with our SCI Inv strain, in which the SCI has become very active thanks to the high folding capacity of all the recombinogenic *attC* strands, including those of the TA cassettes. The growth defect of this strain is also dependent on the integron integrase activity. Indeed, we previously showed that the SCI Inv strain does not have any growth defect in absence of IntIA and that a catalytically active integrase was necessary to induce a growth defect in that strain (Richard et al., 2022b). It seems therefore consistent that events affecting the *intIA* gene, prohibiting its expression or directly affecting its CDS gene, are those most frequently found to attenuate the growth defect.

Concerning the integrase expression, the mutations found in the *araC* regulator of the used pBAD promoter could be considered as an artefact of the experiment. Nevertheless, this confirms that silencing the recombination process abolishes the cost. If, in some genomes, integrase expression is controlled by certain regulators, that could be somewhat representative of this situation. Concerning the mutations affecting the integrase CDS, the most common are nonsense and indels, both of which lead to inactive truncated integrases. Interestingly, half of these mutations have been found in the 5’ part of the integrase gene located in a homopolymer of eight “A”. This type of sequences, defined as short sequence repeat, has already been identified in prokaryotes as hotspots for slipped-strand mispairing (SSM) events during replication (van Belkum et al., 1998). SSM is one of the mechanism responsible for phase variation, an adaptive process that accelerates the rate of spontaneous mutations in bacterial subpopulations, facilitating reversible phenotypic changes and rapid adaptation (Hallet, 2001; Henderson et al., 1999). Here, we provide the first evidence for loss-of-function of an SCI integrase due to SSM events in response to toxicity of unsustainable cassette dynamic. This shows that loss-of-function is the main strategy for bacterial adaptation and that the supply of mutations is the key. We did not see any mutations in the integrase active site, but this is not surprising as there are many ways of inactivating a function without affecting the active site. In addition, the sequences corresponding to the active sites in a protein are generally robust, with only a couple of essential amino acids.

We also demonstrated, by simulating random sequences, that the homopolymers in the SCI integrase genes are not longer than expected, suggesting that there is no selection for their presence. This result is not surprising, since integrase expression, when SCIs are in their natural orientation, does not affect cell growth (Richard et al., 2022b). Interestingly, homopolymers in MI integrases appear to be shorter than expected. This suggests that MIs could be affected by genetic instabilities associated with SSM in their long homopolymers when conjugating within plasmid across species (ssDNA can be more mutagenic). This would lead to a counter-selection of these homopolymers in the MI sequences. The mutational rate due to this type of motifs may thus be slightly lower in MI integrases than in SCI ones. This would be in line with the presumed role of SCI integrases in maintaining a large cassette array, whereas MIs, on the contrary, must be able to efficiently shuffle and widely disseminate their adaptive functions (Escudero et al., 2015; Richard et al., 2022a).

However, the second observed outcome of the evolution experiment – the spontaneous re-inversion of a substantial part (63%) of the SCI cassette array by homologous recombination between the two IS*3* copies – was less expected. Note that we have already observed this type of recombination between repeated IS elements, leading to the fusion of the two chromosomes of *Vibrio cholerae* (Val et al., 2014). However, this is the first time that we observed such a large chromosomal rearrangement inside the *V. cholerae* SCI structure, demonstrating an important role of the duplicated ISs present along the SCIs in their maintenance. We previously showed that the inversion of the SCI led to a higher excision rate of the cassettes by placing the *attC*_*bs*_ on the lagging strand template. Hence, re-inversion of the SCI brings the *attC*_*bs*_ back to their native and less recombinogenic state that is on the leading strand template. The portion of the re-inverted array contains ten non-duplicated TA systems versus only three in the other part (Krin et al., 2023). Indeed, the increased probability of excision of a TA cassette leading to cell death or cell cycle arrest must be a strong selective pressure for the re-inversion of the section of the array that concentrates the most TA cassettes. Finally, the partial re-inversion of the SCI array is another possible means of suppressing the growth defect in the SCI Inv strain expressing the integrase, since it is not the integrase that is intrinsically toxic but the increased excision of the TA cassette. This evolutionary solution has the advantage of keeping integrase functionality intact. Altogether, these results illustrate a tight regulation of the SCI functioning, limiting their evolvability in favour of the stability and cassette maintenance, all orchestrated by a fascinating interplay between gene orientation, bacterial replication, genome rearrangement, toxin–antitoxin systems and bacterial fitness.

## Materials and methods

### Bacterial strains, plasmids and primers

The different bacteria strains, plasmids and primers that were used in this study are described in Table 1, 2 and 3.

**TABLE 1:** Bacterial strains used in this study.

| Strain number | Relevant genotypes or description | References |
| --- | --- | --- |
| **Bacterial strains** |  |  |
| I992, I993, J341 | SCI inverted. *attC<sub>bs</sub>* on lagging strand template (SCI Inv strains) | (Richard et al., 2022b) |
| J053, J054, K025 | SCI re-inverted into its original orientation, *attC<sub>bs</sub>* on leading strand template (SCI Reinv strains) | (Richard et al., 2022b) |
| **Transformed strains** |  |  |
| K483, K485, K487 | SCI Inv + pJ504 | (Richard et al., 2022b) |
| K493, K495, K497 | SCI Reinv + pJ504 | (Richard et al., 2022b) |
| **Evolved strains** |  |  |
| ER332 | K483 culture evolved during 70 generations | This study |
| ER334 | K485 culture evolved during 70 generations | This study |
| ER336 | K487 culture evolved during 70 generations | This study |
| ER338 | K493 culture evolved during 70 generations | This study |
| ER340 | K495 culture evolved during 70 generations | This study |
| ER342 | K497 culture evolved during 70 generations | This study |
| ER229-236 | 8 clones isolated from ER332 | This study |
| ER237-244 | 8 clones isolated from ER334 | This study |
| ER245-252 | 8 clones isolated from ER336 | This study |
| ER253-260 | 8 clones cured from ER229-236 | This study |
| ER261-268 | 8 clones cured from ER237-244 | This study |
| ER269-276 | 8 clones cured from ER245-252 | This study |
| ER277-284 | 8 clones transformed from ER253-260 | This study |
| ER285-292 | 8 clones transformed from ER261-268 | This study |
| ER293-300 | 8 clones transformed from ER269-276 | This study |
| ER179-ER187-ER286 | 3 clones SCI inv with IS3 rec | This study |

**TABLE 2:** Plasmids used in this study.

| Plasmid number | Plasmid description | Relevant properties and construction |
| --- | --- | --- |
| pJ502 | pBAD43$\Delta attC_{aadA1}$ | *oripSC101*; [Sp<sup>R</sup>] (Vit *et al.*, 2021) |
| pJ504 | pBAD43 $\Delta attC_{aadA1}::intIA_{Vch}$ | *oripSC101*; [Sp<sup>R</sup>] (Vit *et al.*, 2021) |

**TABLE 3:**
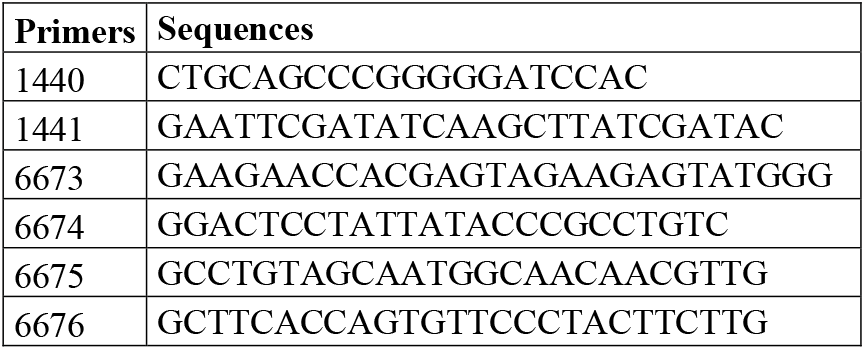
Primers used in this study.

### Media

*Vibrio cholerae* strains were grown in Luria Bertani (LB) at 37°C. Glucose (Glc), L-arabinose (Ara) and fructose (Fruc) were added respectively at final concentrations of 10, 2 and 10 mg/mL. The spectinomycin (Spec) was used at the following concentrations: 100 μg/mL in absence of glucose and 200 µg/mL in presence. To avoid catabolic repression during various tests using arabinose as inducer for the expression of the integrase, cells were grown in a synthetic rich medium: MOPS Rich supplemented with fructose (1%) as a carbon source.

### Evolution experiment

Three clones each stemming from different biological replicates of the SCI Inv and SCI Reinv strains containing the low-copy number pSC101 vector, carrying the *intIA* gene under a pBAD promoter, were grown overnight (O/N) in LB + Spec + Glc. These first cultures correspond to that we called the Day 0 (D0) meaning 0 generation of evolution. To initiate the evolution experiment, each culture was diluted at 1:1000 in 5 ml of the inducing medium (MOPS + Spec + Fruc + Ara). Agitation was set to 180 rpm. Cultures were allowed to grow for 24 h at 37°C (resulting to ∼9.97 generations). This was repeated six times, that is with a total of ∼70 generations (∼10 generations every 24h). Moreover, after 0 (D0), 10 (D1), 20 (D2), 30 (D3), 50 (D5) and 70 (D7) generations, the six independent cultures were streaked on LB agar plates containing Spec + Glc. For the SCI Inv strains with integrase, we isolated 24 independent and randomly chosen clones (8 for each biological triplicates) and for the SCI Reinv strain with integrase, we isolated 8 clones (3, 3 and 2 for each biological triplicates). Each of these clones were grown O/N in LB + Spec + Glc and then used to perform automated growth rate measurements.

### Automated growth rate measurements

O/N cultures of the indicated strains (see above) were diluted 1/1000 in MOPS + Spec + Fruc + Ara medium and then distributed by quadruplicate in 96-well microplates avoiding the use of external rows and columns. Automated growth-curve were generated using a TECAN Infinite microplate reader, with an absorbance measurement of 600 nm taken at 10-min intervals at 37°C on maximum agitation for 15h. Maximum growth rates during exponential phase were directly obtained using the “GrowthRates” R package. The package can be downloaded from CRAN [https://cran.r-project.org/package=growthrates].

### Plasmid curing

In the evolution experiment, the integrase was carried on a pSC101 plasmid and under the control of a pBAD promoter. To replace this plasmid by a new one, we took advantage of the natural instability of plasmids in *V. cholerae* (Jaskolska et al., 2022). 24 isolated clones from the 3 biological replicates evolved after 70 generations were cultivated O/N in LB + Glc (without spec) and then plated on LB agar + Glc plates. The resulting clones were streaked on LB agar + Glc and parallelly on LB agar + Spec + Glc plates. The clones growing on the first but not on the latter where the ones in which the plasmid was cured, which was confirmed by the absence of PCR amplification using the 1440 and 1441 primers. For each of the 24 cured clones obtained, a new plasmid was added by electroporation.

### Electroporation in *V. cholerae*

Cells were grown to late exponential phase (OD_600_ ∼0.7), then 10mL was pelleted. The pellet was washed twice in G buffer (137 mM Sucrose, 1mM HEPES, pH 8.0). 50 uL of the resulting cell suspension was mixed with ∼250 ng of the pSC101::*intIA* (pJ504) plasmid in a 1mm electroporation cuvette. Electroporation was performed at 2000 V. Electroporated cells were grown at 37°C in LB + Glc for 1h for phenotypic expression before plating on LB agar + Spec + Glc plates. The presence of the plasmid was checked by PCR amplification using the o1440 and o1441 primers.

### Plasmid DNA extraction

At the end of the evolution experiment (70 generations), plasmids (containing integrase) of the three independent populations were extracted using the Thermo Fisher Miniprep Kit. The three plasmid preparations were adjusted to 50ng/µL as measured by Qubit, and then mixed at equal proportion prior to Illumina sequencing. The same Kit was used for plasmid extraction from the corresponding isolated clones prior to Sanger sequencing.

### Genomic DNA Extraction

Clones that were to be sequenced or checked by PCR for genome rearrangements were cultivated overnight in LB + Spec + Glc. The genomic DNA was extracted following the specification of the Qiagen gDNA extraction Kit. DNA concentration was assessed by Qubit and quality assessed by migration on a 1% Agar gel prior to sequencing.

### PCR checking of partial SCI re-inversion

Partial SCI re-inversion was checked by PCR using primer pairs i1 (6674 and 6676), i2 (6673 and 6675), r1 (6674 and 6673) and r2 (6676 and 6675). For the partial SCI re-inverted clones, bands are expected only with the r1 and r2 primer pairs (respectively 2380 bps and 2501 bps) and for the non-evolved SCI Inv clones (no partial SCI re-inversion), bands are expected only with i1 and i2 primer couples (2206 bps and 2675 bps respectively). Note that non-evolved SCI inv clones correspond to clones selected from the D0 cultures (obtained in a medium that does not enable integrase expression, i.e. in the presence of glc).

### Sequencing and SNP calling

Both the plasmidic and genomic DNA were sequenced using an Illumina MiSeq sequencer with paired end 150bp reads (2 × 150 bp). The obtained sequenced data were then analysed using the Sequana variant calling pipeline v0.9.5 (Cokelaer et al., 2017), which is a Snakemake-based pipeline. This pipeline involves an initial mapping step performed with the Burrows-Wheeler Alignment software (bwa v0.7.17) (Li and Durbin, 2010) using default parameters. Then, the variant calling was conducted using the Freebayes software v1.3.2 (Garrison and Marth, 2012), which maps the sequencing data and identifies variants. A mutation frequency threshold of 0.1% was applied based on a median coverage of 10,000X along the plasmid’s sequence. Furthermore, the analysis incorporated an assessment of strand balance to validate the detected variants. A tolerance on a strand balance deviating from 0.5 was introduced, and variants with a strand balance exceeding the 0.1 limit were filtered out, contributing to the robustness of the results.

### DNA homopolymer analysis in integron integrase genes

The integrase sequences were extracted from the complete integrons identified with IntegronFinder 2.0 (Neron et al., 2022). We classified them as “mobile” and “sedentary” based on the length of the cassette array, as suggested by the authors. The Nextflow pipeline used to analyze the DNA homopolymers within the extracted integrase genes is available at https://gitlab.pasteur.fr/gmillot/homopolymer and is briefly described here. For each SCI and MI integrase sequence of the input batch: 1) the sequence is split according to homopolymers (e.g., ATTTAACC is split into A, TTT, AA, CC), 2) homopolymer lengths were enumerated, independently of nucleotide content, which returns the observed counting and related proportions of homopolymer lengths in the sequences, 3) the two first steps were re-applied after randomly shuffling the nucleotides of each sequence, 4) This was performed 10,000 times and means of homopolymer lengths were computed, which returns the theoretical counting and related proportions of homopolymer lengths in the sequence. All the homopolymer results are available at https://doi.org/10.5281/zenodo.8305871.

## Data availability

All data are available in the main text or the supplementary materials. All plasmid and genomic sequences are publicly available in The European Nucleotide Archive (ENA). The accession no. for the genomic sequences of the 3 clones SCI inv with IS*3* rec are ERS16316977 (ER179), ERS16316978 (ER187) and ERS16316979 (ER286). The accession no. for the plasmid sequences are ERS16316976 (pJ504_evo_SCI inv) and ERS16446759 (pJ504_evo_SCI reinv). All other raw data are available upon request to the corresponding author.

## Acknowledgements

We thank Marc Monot, Laurence Ma, Juliana Pipoli Da Fonseca and Laure Lemée from the Biomics platform, C2RT, Institut Pasteur, Paris, France, supported by France Génomique (ANR-10-INBS-09) and IBISA.

